# Hemodynamic responses link individual differences in informational masking to the vicinity of superior temporal gyrus

**DOI:** 10.1101/2020.08.21.261222

**Authors:** Min Zhang, Nima Alamatsaz, Antje Ihlefeld

**Author notes:** **For correspondence** (AI). **Present address:** Department of Biomedical Engineering, New Jersey Institute of Technology, Newark NJ, USA.

## Abstract

Suppressing unwanted background sound is crucial for aural communication. A particularly disruptive type of background sound, informational masking (IM), often interferes in social settings. However, IM mechanisms are incompletely understood. At present, IM is identified operationally: when a target should be audible, based on suprathreshold target/masker energy ratios, yet cannot be heard because perceptually similar background sound interferes. Here, functional near infrared spectroscopy recordings show that task-evoked blood oxygenation changes near the superior temporal gyrus (STG) covary with behavioral speech detection performance for high-IM but not low-IM background sound, suggesting that the STG is part of an IM-dependent network. Moreover, listeners who are more vulnerable to IM show increased hemodynamic recruitment near STG. In contrast, task-evoked responses near another auditory region of cortex, the caudal inferior frontal sulcus (cIFS), do not predict behavioral sensitivity, suggesting that the cIFS belongs to an IM-independent network. Results are consistent with the idea that cortical gating shapes individual vulnerability to IM.

## Introduction

Perceptual interference from background sound, also called auditory masking, has long been known to impair the recognition of aurally presented speech through a combination of at least two mechanisms. Energetic masking (EM) occurs when target and masker have energy at the same time and frequency, such that the masker swamps or suppresses the auditory nerve activity evoked by the target (***Young and Barta, 1986***; ***Delgutte, 1990***). Informational masking (IM) is presently defined operationally. IM occurs when a target is expected to be audible based on EM mechanisms, yet cannot be detected or identified. Listeners experience IM when target and masker are perceptually similar to each other (e.g., hearing two women talk at the same time *vs* hearing out a female in the background of a male voice; ***Brungart*** (***2001b***)) or when the listener is uncertain about perceptual features of the target or masker (e.g., trying to hear out a target with known *vs* unexpected temporal patterning, cf. ***Lutfi et al.*** (***2013***)).

Unlike EM, IM is associated with striking variation in individual vulnerability (***Neff and Dethlefs, 1995***; ***Durlach et al., 2003***). Moreover, an individual’s susceptibility to IM is largely refractory to training (***Neff et al., 1993***; ***Oxenham et al., 2003***). Identifying brain regions where IM-evoked activation patterns covary with individual differences in behavioral vulnerability to IM may thus hold a key for defining the neural mechanisms underlying IM.

Neuroimaging studies have greatly advanced our understanding of the neural mechanisms of masking. Converging evidence links both EM and IM to recruitment of superior temporal gyrus (STG) and frontal cortex (***Davis and Johnsrude, 2003***, ***2007***; ***Scott et al., 2004***, ***2006***, ***2009***; ***Mesgarani and Chang, 2012***; ***Lee et al., 2013***; ***Michalka et al., 2015***). For instance, the predominantly activated STG hemisphere can shift depending on the amount of IM in the background sound (***Scott et al., 2009***). Moreover, for speech that was either spectrally degraded or had impoverished amplitude cues, spanning the range from unintelligible to fully intelligible, activation near STG can account for approximately 40 to 50% of the variance in speech intelligibility (***Pollonini et al., 2014***; ***Lawrence et al., 2018***).

In addition, lateral frontal cortex engages more strongly with increasing listening effort or increasing recruitment of higher-order semantic processes (***Davis and Johnsrude, 2003***; ***Scott et al., 2004***; ***Wild et al., 2012***; ***Wijayasiri et al., 2017***). Parts of lateral frontal cortex, including the caudal inferior frontal sulcus (cIFS), are also sensitive to auditory short-term memory load in situations with IM (***Michalka et al., 2015***; ***Noyce et al., 2017***). Using functional near-infrared spectroscopy (fNIRS), we previously confirmed that the cIFS region engages more strongly when listeners actively attend to speech in IM *vs* listen passively (***Zhang et al., 2018***), making the STG and cIFS promising region of interest (ROIs) for the current study.

Widening an established IM paradigm (***Arbogast et al., 2002***), we here compare hemodynamic responses to low *vs* high IM speech. We test two hypotheses. H1: Individual differences in vulnerability to IM are mediated through processing limitations in the vicinity of STG. H2: Individual differences in vulnerability to IM arise near cIFS.

To study how cortical responses shape individual differences in behavioral speech comprehension, our goal is to differentiate between brain areas with IM independence (task-evoked responses do not predict vulnerability to IM) *vs* areas with IM dependence (task-evoked responses predict IM vulnerability). Using psychometric testing and fNIRS, we simultaneously quantify behavioral sensitivity and hemodynamic responses in the vicinity of STG and cIFS. In experiment 1, we contrast hemodynamic responses to speech detection in presence of combined low-IM *vs* high-IM with same-ear masking. To control for EM, in experiment 2, we contrast high-IM with same-ear *vs* opposite-ear masking. The two experiments serve as their own control, confirming test-retest reliability of the measured cortical traces. Our results support H1 but not H2.

## Results

### Experiment 1

Using the setup shown in Figure 1A, we recorded hemodynamic responses near cIFS and STG bilaterally, from normal-hearing young individuals. Listeners were instructed to detect when the target voice on the left uttered color keywords while SPEECH *vs* NOISE maskers interfered from the right side (Figure 1B). Behavioral pilot testing confirmed that these spectrally sparse maskers produced high-IM (SPEECH) *vs* low-IM (NOISE, Supplement 1).

**Figure 1.**
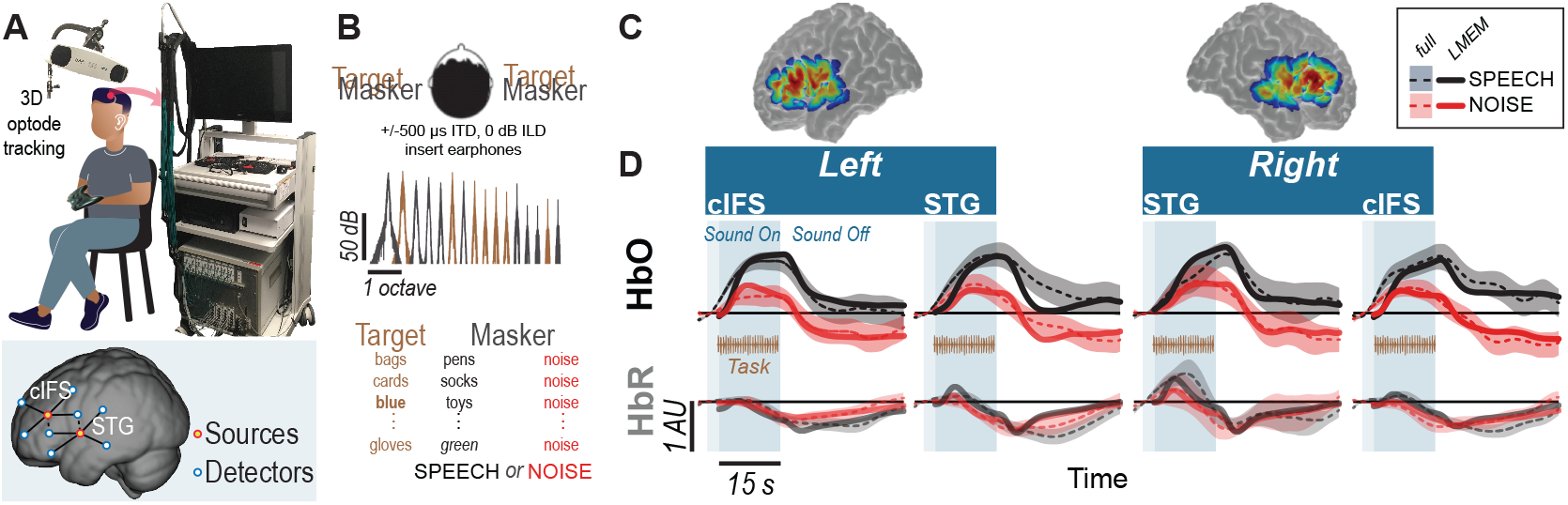
High-IM elicits stronger task-evoked responses than low-IM across all tested ROIs in experiment 1. (A) Experimental apparatus and setup and optode placement for a representative listener. Blue circles show placements of detector optodes, red circles of source optodes (deep channels: solid lines; reference channels: dashed lines). (B) Task design for SPEECH *vs* NOISE. Both target (left-leading ITD of −500 *μs*) and masker (right-leading ITD of 500 *μs*) were presented binaurally. Spectral densities for target *vs* masker show mutually flanking, sharply tuned component bands. (C) Sensitivity maps for optodes placed in the vicinity of STG and cIFS. Warmer colors denote increased likelihood that photons will be recorded from these areas. (D) HbO (top) and HbR (bottom) traces. Full hemodynamic responses are denoted by solid lines and error ribbons. Here and elsewhere, ribbons show one standard error of the mean across listeners. Task-evoked hemodynamic responses predicted from the linear mixed effects model (LMEM) are shown as dashed lines. Shaded areas mark the task duration.

Accounting for approximately half of the variance in the recorded traces (*R*^2^ = 0.45), a Linear Mixed Effects Model (LMEM) was then used to predict task-evoked hemodynamic responses, by regressing out reference channels (*β*_6_ and *β*_7_), block number (*β*_5_), and PTA (*β*_11_ and *β*_12_) from the full response (Supplement 2). Note that the reference channels comprise 44.6% of the total activation levels in the LMEM fits, as calculated via the area under the fitted curve with *vs* without *β*_6_ and *β*_7_. Indeed, unlike the full hemodynamic response, the LMEM-estimated task-evoked hemodynamic response aligns well with the task-onset (compare onset of darker shaded area and dashed line throughout 1D).

LMEM fits reveal significant task-evoked responses at all four ROIs (Table 1; *β*_1−4_ > 0, *p* < 0.0001; see Figure 1D for HbO (top row) and HbR traces (bottom row). Moreover, all ROIs were sensitive to IM. Activation was stronger in the SPEECH as compared to the NOISE configuration (*β*_10_ > 0). The size of the difference between SPEECH (black lines in Figure 1D) *vs* NOISE (red lines) activation varied across ROIs, but these interactions with ROI were small compared to the overall effect size(interaction between masker configuration and cortical structure: *β*_13_ < 0; interaction between masker configuration and hemisphere: *β*_14_ < 0; see Supplement 3).

**Table 1.** Results of LMEM, experiment 1.

|  | Term | Estimate | S.E. | t | p |  |
| --- | --- | --- | --- | --- | --- | --- |
| $\beta_0$ | Intercept | -0.35 | 0.092 | -3.8 | 0.0001 | *** |
| $\beta_1$ | HRF<sub>HbO</sub> | 0.55 | 0.004 | 138.3 | <0.0001 | *** |
| $\beta_2$ | HRF'<sub>HbO</sub> | 0.17 | 0.004 | 39.4 | <0.0001 | *** |
| $\beta_3$ | HRF<sub>HbR</sub> | 0.02 | 0.004 | 5.8 | <0.0001 | *** |
| $\beta_4$ | HRF'<sub>HbR</sub> | 0.11 | 0.043 | 26.8 | <0.0001 | *** |
| $\beta_5$ | Block Number | 0.01 | 0.000 | 76.6 | <0.0001 | *** |
| $\beta_6$ | Reference Channel<sub>HbO</sub> | 0.42 | 0.000 | 1342.0 | <0.0001 | *** |
| $\beta_7$ | Reference Channel<sub>HbR</sub> | 0.44 | 0.001 | 580.8 | <0.0001 | *** |
| $\beta_8$ | Hemisphere | 0.04 | 0.028 | 1.5 | 0.14 | |
| $\beta_9$ | Cortical Structure | 0.08 | 0.026 | 3.0 | 0.003 | ** |
| $\beta_{10}$ | Masker | 0.14 | 0.061 | 2.2 | 0.025 | * |
| $\beta_{11}$ | R ear PTA | 0.02 | 0.008 | 1.8 | 0.08 | . |
| $\beta_{12}$ | L ear PTA | -0.01 | 0.005 | -0.9 | 0.38 | |
| $\beta_{13}$ | Masker Configuration : Cortical Structure | -0.03 | 0.003 | -12.8 | <0.0001 | *** |
| $\beta_{14}$ | Masker Configuration : Hemisphere | -0.05 | 0.003 | -21.1 | <0.0001 | *** |
| $\beta_{15}$ | Cortical Structure : Hemisphere | -0.01 | 0.003 | -5.4 | <0.0001 | *** |
| $\beta_{16}$ | HRF<sub>HbO</sub> : Masker Configuration | -0.19 | 0.004 | -46.5 | <0.0001 | *** |
| $\beta_{17}$ | HRF<sub>HbO</sub> : Cortical Structure | 0.17 | 0.004 | 41.6 | <0.0001 | *** |
| $\beta_{18}$ | HRF<sub>HbO</sub> : Hemisphere | -0.43 | 0.004 | -10.8 | <0.0001 | *** |
| $\beta_{19}$ | HRF'<sub>HbO</sub> : Masker Configuration | 0.02 | 0.004 | 5.6 | <0.0001 | *** |
| $\beta_{20}$ | HRF'<sub>HbO</sub> : Cortical Structure | -0.22 | 0.004 | -51.6 | <0.0001 | *** |
| $\beta_{21}$ | HRF'<sub>HbO</sub> : Hemisphere | -0.04 | 0.004 | -9.5 | <0.0001 | *** |
| $\beta_{22}$ | HRF<sub>HbR</sub> : Masker Configuration | -0.12 | 0.004 | -30.2 | <0.0001 | *** |
| $\beta_{23}$ | HRF<sub>HbR</sub> : Cortical Structure | -0.01 | 0.004 | -1.0 | 0.3 | |
| $\beta_{24}$ | HRF<sub>HbR</sub> : Hemisphere | 0.05 | 0.004 | 11.9 | <0.0001 | *** |
| $\beta_{25}$ | HRF'<sub>HbR</sub> : Masker Configuration | -0.10 | 0.004 | -22.4 | <0.0001 | *** |
| $\beta_{26}$ | HRF'<sub>HbR</sub> : Cortical Structure | 0.16 | 0.004 | 36.6 | <0.0001 | *** |
| $\beta_{27}$ | HRF'<sub>HbR</sub> : Hemisphere | 0.04 | 0.004 | 9.3 | <0.0001 | *** |

Source: Link\_to\_raw\_data **Table 1-source data 1.**
All estimates are referenced to a default condition in left cIFS for SPEECH.
Significance codes: '\*\*\*' $p < 0.001$ , '\*\*' $p < 0.01$ , '\*' $p < 0.05$ , '.' $p < 0.1$ , ' ' $p > 0.1$ . Note: "Int" = "intercept"; "S.E." = standard error of the mean

**Table 2.** Results of LMEM, experiment 2.

|  | Term | Estimate | S.E. | t | p |  |
| --- | --- | --- | --- | --- | --- | --- |
| $\beta_0$ | Intercept | 0.02 | 0.065 | 0.3 | 0.75 | |
| $\beta_1$ | HRF<sub>HbO</sub> | 0.29 | 0.003 | 89.4 | <0.0001 | *** |
| $\beta_2$ | HRF'<sub>HbO</sub> | 0.07 | 0.003 | 19.5 | <0.0001 | *** |
| $\beta_3$ | HRF<sub>HbR</sub> | -0.04 | 0.003 | -10.9 | <0.0001 | *** |
| $\beta_4$ | HRF'<sub>HbR</sub> | 0.07 | 0.004 | 20.8 | <0.0001 | *** |
| $\beta_5$ | Block Number | 0.01 | 0.000 | 39.1 | <0.0001 | *** |
| $\beta_6$ | Reference Channel<sub>HbO</sub> | 0.67 | 0.001 | 1490.4 | <0.0001 | *** |
| $\beta_7$ | Reference Channel<sub>HbR</sub> | 0.73 | 0.001 | 802.0 | <0.0001 | *** |
| $\beta_8$ | Hemisphere | -0.02 | 0.025 | -0.7 | 0.46 | |
| $\beta_9$ | Cortical Structure | 0.04 | 0.034 | 1.2 | 0.23 | |
| $\beta_{10}$ | Masker | 0.00 | 0.025 | 0.04 | 0.97 | |
| $\beta_{11}$ | R ear PTA | -0.01 | 0.011 | -0.97 | 0.33 | |
| $\beta_{12}$ | L ear PTA | 0.00 | 0.009 | 0.3 | 0.79 | |
| $\beta_{13}$ | Masker Configuration : Cortical Structure | 0.06 | 0.002 | 26.3 | <0.0001 | *** |
| $\beta_{14}$ | Masker Configuration : Hemisphere | -0.03 | 0.00 | -14.5 | <0.0001 | *** |
| $\beta_{15}$ | Cortical Structure : Hemisphere | 0.08 | 0.002 | 40.3 | <0.0001 | *** |
| $\beta_{16}$ | HRF<sub>HbO</sub> : Masker Configuration | -0.1 | 0.003 | -31.8 | <0.0001 | *** |
| $\beta_{17}$ | HRF<sub>HbO</sub> : Cortical Structure | 0.04 | 0.003 | 11.1 | <0.0001 | *** |
| $\beta_{18}$ | HRF<sub>HbO</sub> : Hemisphere | 0.03 | 0.003 | 8.5 | <0.0001 | *** |
| $\beta_{19}$ | HRF'<sub>HbO</sub> : Masker Configuration | -0.01 | 0.003 | -1.8 | 0.072 | . |
| $\beta_{20}$ | HRF'<sub>HbO</sub> : Cortical Structure | -0.19 | 0.003 | -53.9 | <0.0001 | *** |
| $\beta_{21}$ | HRF'<sub>HbO</sub> : Hemisphere | -0.06 | 0.003 | -16.63 | <0.0001 | *** |
| $\beta_{22}$ | HRF<sub>HbR</sub> : Masker Configuration | 0.003 | 0.003 | 1.1 | 0.29 | |
| $\beta_{23}$ | HRF<sub>HbR</sub> : Cortical Structure | -0.05 | 0.003 | -14.4 | <0.0001 | *** |
| $\beta_{24}$ | HRF<sub>HbR</sub> : Hemisphere | -0.04 | 0.003 | -11.9 | <0.0001 | *** |
| $\beta_{25}$ | HRF'<sub>HbR</sub> : Masker Configuration | 0.01 | 0.003 | 3.0 | 0.0031 | ** |
| $\beta_{26}$ | HRF'<sub>HbR</sub> : Cortical Structure | 0.06 | 0.003 | 17.8 | <0.0001 | *** |
| $\beta_{27}$ | HRF'<sub>HbR</sub> : Hemisphere | -0.01 | 0.003 | -3.5 | 0.0006 | *** |

Source: Link\_to\_raw\_data Table 2-source data 1.
All estimates are referenced to a default condition in left cIFS for SPEECH. Significance codes: '\*\*\*' p < 0.001,
'\*\*' p < 0.01, '\*' p < 0.05, '.' p < 0.1, '' p > 0.1. Note: "Int" = "intercept"; "S.E." = standard error of the mean

### Experiment 2

The sharply tuned, mutually flanking bands of target and masker in experiment 1 were presented to both ears, and were designed to produce high- *vs* low IM, with little EM. However, IM can also occur when target and masker are presented to opposite ears. It is unclear whether the neural mechanisms underlying IM are similar when target and masker are presented to the same *vs* opposite ears. Thus, we next wished to examine whether the pattern of STG and cIFS recruitment would generalize to a dichotic IM configuration.

Testing a new group of 14 listeners, experiment 2 contrasted SPEECH with SPEECH-oppo, a stimulus configuration that was identical to SPEECH, except that target and masker were now presented to opposite ears (Figure 2). Mirroring results from experiment 1, an LMEM fitting all HbO and HbR traces from experiment 2 accounted for approximately half of the variance in the recorded data (*R*^2^ = 0.52), with 60.2% of the full hemodynamic activation attributed to reference channels. Moreover, LMEM fits confirmed that task-evoked responses in all four ROIs occurred in both masker configurations, even when target and masker were presented to opposite ears (2;*β*_1−4_ > 0, *p* < 0.0001). All ROIs engaged more strongly in the SPEECH as compared to the SPEECH-oppo configuration (*β*_10_ > 0), with effect size depending somewhat on ROI (see Supplement 3).

**Figure 2.**
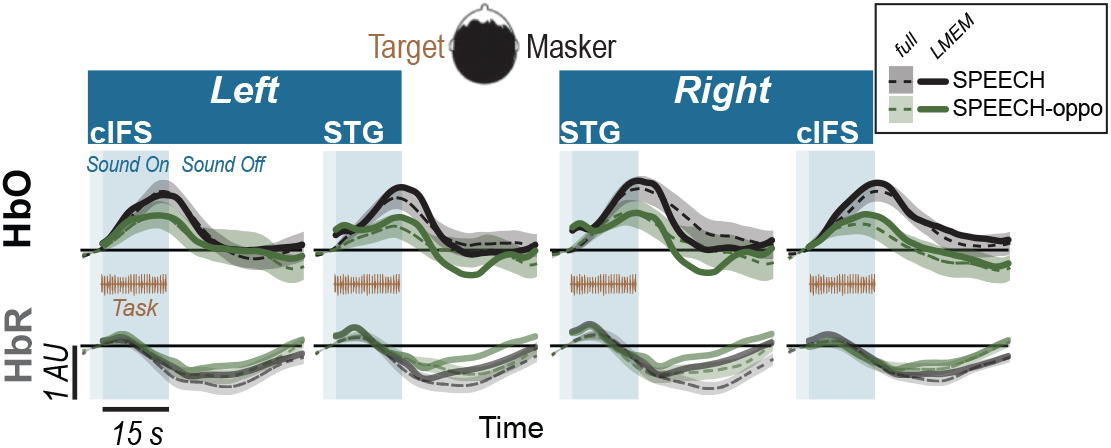
Hemodynamic responses for SPEECH (Black) *vs* SPEECH-oppo (green) show robust task-evoked recruitment of all ROIs in experiment 2, even when target and masker are presented to opposite ears. Solid lines and error ribbons denote raw recordings; dashed lines show LMEM fits.

### Vulnerability to masking and hemodynamic responses

To test the core hypotheses, we next examined STG and cIFS for IM-dependence. We reasoned that in an IM-dependent ROI, the hemodynamic activation strength should predict behavioral sensitivity.

For each ROI, planned adjusted coefficients of determination, *R*^2^, between behavioral speech detection sensitivity and the peak of the HbO response were calculated. In experiment 1, individual behavioral thresholds were significantly anti-correlated with peak HbO only in the SPEECH configuration in the vicinity of left or right STG, where hemodynamic responses explained 23% (left STG) and 31% (right STG) of the behavioral variance (black square symbols in Figure 3A). In contrast, behavioral NOISE thresholds were uncorrelated with hemodynamic responses (Figure 3B). Note that these differences in hemodynamic activation patterns were observed despite the fact that the behavioral speech detection performance, measured during the fNIRS recordings, was comparable between NOISE and SPEECH [paired t-test: *t*(13) = −1.14, *p* = 0.27]. Furthermore, activity levels near cIFS (Figure 1C) were not correlated with behavioral thresholds in SPEECH or NOISE.

**Figure 3.**
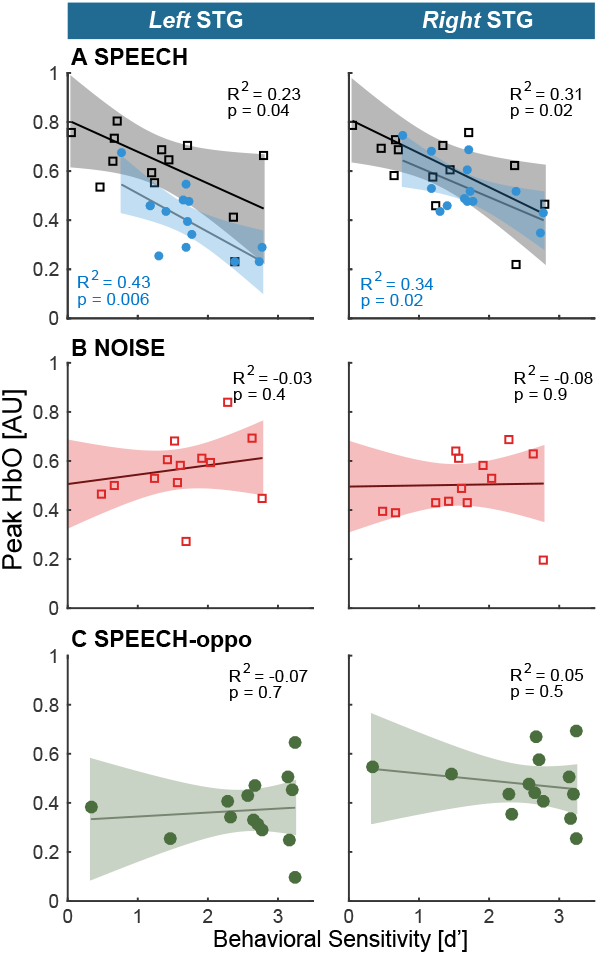
Hemodynamic responses link individual differences in vulnerability towards IM to the vicinity of STG. (A) STG activity and behavioral vulnerability to the high-IM SPEECH condition are robustly anti-correlated, across both hemispheres in experiments 1 and 2 (black *vs* blue symbols, respectively). (B) There was no appreciable association between HbO peaks and the low-IM NOISE condition. (C) When target and masker were presented to opposite ears in the SPEECH-oppo configuration, HbO peaks did not predict psychophysical thresholds.

Testing a different group of listeners, experiment 2 confirmed the finding from experiment 1 that HbO peaks near left or right STG were significantly anti-correlated with behavioral sensitivity for the SPEECH configuration. Moreover, activity levels in cIFS were again uncorrelated with behavioral thresholds. Identical SPEECH configurations were assessed in experiments 1 and 2. Therefore, the converging results across two groups of listeners confirm high test-retest reliability of the current fNIRS approach. Specifically, in experiment 2, STG HbO peak activation explained 43% and 34% of the behavioral variance in left and right STG respectively (blue square symbols in Figure 3A). In contrast, hemodynamic responses for SPEECH-oppo did not predict behavioral sensitivity (Figure 3C).

A caveat, unlike in experiment 1, in experiment 2, task difficulty differed across masking conditions. Specifically, behavioral speech detection thresholds were better for SPEECH-oppo than SPEECH [paired t-test: *t*(13) = −3.13, *p* = 0.008; compare green symbols in Figure 3C falling to the right of the red, blue and black symbols in Figure 3A,B]. However, even for the more poorly performing listeners in experiment 2, no obvious trend links behavioral sensitivity to peak HbO levels in left or right STG.

Of note, behavioral responses were not predicted from HbR activity levels, across any of the tested conditions, in either of the two experiments. As expected, task-evoked HbO and HbR responses were robustly anti-correlated (in Figures 1D, and 2, compare dark dashed lines in the top row to the lighter dashed lines of the same color in the bottom row). This anti-correlation would predict that HbR responses mirror the correlation patterns between HbO peaks and behavioral sensitivity. However, in general, HbR response magnitudes were very small, approximately 20% of HbO magnitudes, hinting that here, the HbR responses may have been contaminated by the noise floor of the recording system.

## Discussion

The goal of the current work was to identify brain regions where individual differences in IM vulnerability emerge. To that end, we sought to differentiate between IM-independent parts of the brain whose activation levels are equivalently driven by low- or high-IM, *vs* IM-dependent regions whose activation levels correlate with individual IM-vulnerability.

### Hemodynamic correlates of IM

The current data confirm that cortical regions at or near STG and cIFS engage during masked speech comprehension tasks (***Scott et al., 2004***, ***2006***, ***2009***; ***Rowland et al., 2018***; ***Kerlin et al., 2010***; ***Mesgarani and Chang, 2012***; ***Ding and Simon, 2012***; ***Michalka et al., 2015***; ***Noyce et al., 2017***; ***Zhang et al., 2018***). For both high- and low-IM background sound, when a listener engaged in speech detection, robust task-evoked hemodynamic responses in STG and cIFS occurred in both brain hemispheres. Task-evoked bilateral responses in STG and cIFS were even observed when target and high-IM masker were presented to opposite ears (SPEECH-oppo in experiment 2).

SPEECH masking recruited a stronger task-evoked response than NOISE masking in both left and right STG, consistent with prior work (***Scott et al., 2004***). Activation levels during SPEECH masking consistently predicted 20-43% of individual differences in vulnerability in left or right STG, in both experiments. Moreover, STG recruitment did not predict vulnerability to masking for the low-IM masker (NOISE condition in experiment 1). Together, these results show that recruitment in the vicinity of STG was IM-dependent. In contrast, while cIFS also showed task-evoked responses that were stronger in SPEECH than in NOISE, cIFS activation strength did not significantly correlate with individual vulnerability in any tested masking configuration, suggesting that the vicinity of cIFS was IM-independent.

IM is thought to be a central auditory mechanism. However, IM generally interferes much more strongly when target and masker are presented to the same ear(s), as compared to being presented to opposite ears (***Brungart and Simpson, 2002***, ***2007***; ***Kidd Jr et al., 2003***; ***Gallun et al., 2005***; ***Wightman and Kistler, 2005***). It is unclear whether these mechanisms are similar for same-ear *vs* opposite ear IM. Even when background sound enters a non-target ear, behavioral evidence suggests that IM interference can be attributed to a combination of a failure to attend to the target ear as well as increased listening effort (***Gallun et al., 2007***), whereas same-ear masking adds the possibility that energetic masking shapes IM through interactions with attention and across-time streaming (***Ihlefeld and Shinn-Cunningham, 2008***).

Here, SPEECH-oppo evoked bilateral responses in STG and cIFS. If identical STG-based networks were activated for same-ear-IM (SPEECH) and opposite-ear-IM (SPEECH-oppo), STG activity should have been a negative predictor of behavioral SPEECH-oppo sensitivity, but this was not observed. A caveat, speech identification thresholds in SPEECH-oppo were close to ceiling for a few of the listeners. However, even for poorly performing listeners, no trend emerged linking the peak HbO response and behavioral sensitivity (Figure 3C). Moreover, the interpretation that contralateral IM recruits different brain networks than ipsilateral IM is also supported by prior evidence from research in children, where the ability to suppress a masker ipsilateral to the target matures more slowly than the ability to suppress a masker on the contralateral side (***Wightman et al., 2010***).

For same-ear IM, listeners reached comparable speech detection thresholds in low-IM and high-IM, but had marked individual difference during IM speech identification during behavioral pilot testing. This observation is consistent with the idea that more IM-vulnerable listeners exerted more listening effort (***Pichora-Fuller et al., 2016***). A cortical marker for listening effort was previously located in lateral inferior frontal gyrus, a brain area which shows attention-dependent increase in frontal brain activation during listening to degraded speech (***Wild et al., 2012***; ***Wijayasiri et al., 2017***). The current study did not target the lateral inferior frontal gyrus, nor did we record alternative measures of listening effort, such as pupilometry (***Zekveld and Kramer, 2014***; ***Parthasarathy et al., 2020***), precluding any direct test of this possibility.

Together, the results show that even with comparable behavioral sensitivities and similar long-term acoustic energy, high-IM in the same ear increased HbO peaks near STG and cIFS, as compared to low-IM. This effect was observed separately for same-ear as well as opposite-ear IM. Moreover, the observed anti-correlation between HbO peak levels and individual task performance in same-ear high-IM is consistent with the interpretation that left and right STG are part of a same-ear-IM-dependent network. In contrast, the vicinity of cIFS engaged in an IM-independent manner.

### Emergence of IM

Listeners with higher cognitive abilities comprehend masked speech better (***Mattys et al., 2012***; ***Rönnberg et al., 2008***), but prior work shows no evidence that cognitive ability contributes differently to IM *vs* EM. For instance, cognitive scores poorly predict how well an individual can utilize an auditory scene analysis cue to suppress IM (***Füllgrabe et al., 2015***). Consistent with this, here, task-evoked responses near cIFS were IM-independent, unlike in the vicinity of STG.

Inded, prior work hints that IM emerges at the level of auditory cortex, a part of the STG (***Gutschalk et al., 2008***). We here tested maskers that were spectrally interleaved with the target, designed to produce either high IM (SPEECH) or low IM (NOISE). EM, when present, was limited to spectral regions outside the frequency bands that comprised most of the target energy. Consistent with this, for speech *detection*, behavioral thresholds were comparable between SPEECH and NOISE. However, our behavioral pilot results also confirmed that speech *identification* was much more difficult in the presence of SPEECH than NOISE (***Freyman et al., 1999***; ***Arbogast et al., 2002***; ***Brungart et al., 2006***; ***Wightman et al., 2010***).

This behavioral pattern parallels a behavioral phenomenon in vision - called Crowding. In Crowding, the presence of visual target identification is severely impaired by nearby clutter or “flankers” (***Bouma, 1970***; ***Rosen et al., 2014***). In the current IM design, the spectrally sparse masker and target can be conceptualized as mutually flanking each other. Moreover, analogous to the current behavioral results, flankers that Crowd target identification do not affect target detection (***Pelli et al., 2001***). Furthermore, using a behavioral paradigm that is comparable to the current speech identification task, prior work shows that IM can occur even when the masker is softer than the target (***Brungart, 2001a***; ***Ihlefeld and Shinn-Cunningham, 2008***). Analogously, Crowding can occur even when the flankers are smaller than the target (***Pelli et al., 2001***). Of importance to the current work, there is good evidence that the Crowding effect occurs in the visual cortex (***Millin et al., 2014***; ***Zhou et al., 2018a***). In particular, flankers presented through one eye crowd a target presented through the other eye (***Flom et al., 1963***; ***Taylor and Brown, 1972***; ***Tripathy and Levi, 1994***). These striking similarities of IM and Crowding suggest that they result from analogous sensory processes, further supporting the prior notion that IM arises at the level of cortex.

### Cortical mechanisms of IM

The current results show that for similar behavioral sensitivities and similar long-term acoustic energy, individual differences in vulnerability to high-IM in the same ear correlated with increased need for supply of oxygen in the vicinity of STG, as compared to low-IM. However, converging evidence from prior work with electroencephalography (EEG) recordings also shows that the temporal fidelity by which cortical local field potentials encode sound, as opposed to their absolute response strength, correlates with task demands and predicts masked speech intelligibility (***Choi et al., 2014***; ***O’Sullivan et al., 2015***; ***Viswanathan et al., 2019***). Note that unlike with hemodynamic responses recorded with fNIRS, which emerge within proximity of the recording sensors at STG, it is generally more difficult to pinpoint where in the brain the EEG traces originate. In addition, even listeners with audiologically normally hearing can vary dramatically in their ability to resolve and utilize temporal fine structure cues (***Ruggles et al., 2011***; ***Bharadwaj et al., 2019***). Moreover, an individual’s sensitivity to monaural or binaural temporal fine structure predicts masked speech intelligibility, especially in temporally fluctuating background sound (***Lorenzi et al., 2006***; ***Papesh et al., 2017***). Intriguingly, the neural mechanisms shaping temporal fidelity are thought to be of *sub*cortical origin (***Parthasarathy et al., 2020***). Furthermore, prior work with MEG indicates that a thalamo-cortical loop gates temporal signatures of sound to the cortical processing level (***Bharadwaj et al., 2016***). Consistent with this, recent cortical recordings in humans also demonstrate that neural tuning properties of the STG rapidly and flexibly shift in gain, temporal sensitivy and spectrotemporal tuning, depending on the stimulus (***Khalighinejad et al., 2019***; ***Keshishian et al., 2020***).

Together, these findings raise the possibility that an individual’s need for gating or adapting the neural code in STG should increase with decreasing temporal fidelity of subcortical information, as they need to work harder to overcome poor subcortical encoding of the target. Increased inhibitory activity in STG associated with stronger modulation or gating of subcortical temporal fidelity in vulnerable listeners should therefore increase the amplitude of hemodynamic responses (***Stefanovic et al., 2004***; ***Vazquez et al., 2018***). Broadly increased inhibition would not necessarily be picked up via EEG analysis looking for temporal coherence and/or EEG recordings summing neural activity farther from STG. Thus, the current results are consistent the idea that increased gating or modulation of subcortical information via STG may be a potential mechanism contributing for individual variability in IM vulnerability. Future work is needed to explore how metabolic need and the fidelity of cortical temporal coding interact.

### Spatial Specificity

The spacing of fNIRS optodes determines both the depth of the brain where recorded traces originate, as well as their spatial resolution along the surface of the skull. Here, optode sources and detectors were spaced 3 cm apart and arranged cross-wise around the center of each ROI (Figure 1A). To estimate the hemodynamic activity in each ROI, we averaged across the four channels of each ROI. This averaging greatly improved test-retest reliability of each ROI’s activation trace during pilot testing, both here and in our prior work (***Zhang et al., 2018***). A caveat of this approach is that it reduces the spatial resolution of the recordings. Thus, it is unclear whether increased hemodynamic activity near STG is due to increased STG recruitment, or due to a more broadly activated brain network in the vicinity of STG. For instance, there is precedence for activation of additional brain regions as a compensatory strategy for coping with age-related cognitive decline (***Presacco et al., 2016***; ***Jamadar, 2020***). Listeners who are more vulnerable may use either a broadened brain network or increase STG recruitment, two possibilities that the current data cannot differentiate. However, either interpretations is consistent with the idea that a central processing limitation exists that includes STG and shapes vulnerability to IM.

### Diagnostic Utility

The current results bear clinical relevance. A technique we here used to design our stimuli, vocoding, is a core principle of speech processing with current cochlear implants. A pressing issue for the majority of cochlear implant users is that they cannot hear well in situations with masking, an impairment in part attributed to cortical dysfunction (***Anderson et al., 2017***; ***Zhou et al., 2018b***). Sending target and masker sound to opposite ears can improve target speech identification in some, but not all, bilateral cochlear implant users of comparable etiology, suggesting that central auditory processing contributes to clinical performance outcomes (***Goupell et al., 2016***). However, a challenge for imaging central auditory function in cochlear implant users is that cochlear implants are ferromagnetic devices. Thus, cochlear implants often either unsafe for use in magnetic resonance imaging (MRI) scanners and/or cause sizeable artifacts when imaged with MRI or EEG (***Hofmann and Wouters, 2010***). Moreover, when imaged under anesthesia, cochlear implant stimulation can fail to elicit cortical responses (***Nourski et al., 2013***). In contrast, fNIRS, a quiet and light-based technology, is safe to use with cochlear implants. The current paradigm demonstrates that fNIRS-recorded cortical responses to masked speech with impoverished, cochlear-implant-like qualities, can explain approximately a third of the variance in individual vulnerability to IM - an approach that, it is hoped, may prove useful in future clinical practice.

## Methods and Materials

### Participants

Our sample size (14 participants for each of the two fNIRS experiments and 11 participants for a behavioral pilot control) was selected *a priori* using effect size estimates from prior work on IM (***Zhang et al., 2018***; ***Arbogast et al., 2002***). In total, we recruited 40 paid listeners, who were right-handed native speakers of English, and between 19 and 25 years old (17 females). Assessment of pure-tone audiometric detection thresholds (PTAs) at all octave frequencies from 250 Hz to 8 kHz of 20 dB HL or better verified that all listeners had normal hearing. Specifically, the across-ear differences in pure tone thresholds was 10 dB or less, at all of the audiometric frequencies. All listeners gave written informed consent prior to participating in the study. All testing was administered according to the guidelines of the Institutional Review Board of the New Jersey Institute of Technology.

### Speech Stimuli

There were 16 possible English words, each utterance recorded without co-articulation by each of two male talkers (Kidd, et al. 2008). The words consisted of the colors <red, white, blue, and green> and the objects <hats, bags, cards, chairs, desks, gloves, pens, shoes, socks, spoons, tables, and toys>. The colors were designated as keywords. Target word sequences were generated by picking a total of 25 random words from the overall set of 16, including between three and five target words, and concatenating them in random order with replacement (a set of more than 10^26^ possible permutations for the target sequence, 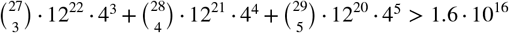). Similarly, masker sequences were made by picking 25 random words from the overall set of 16, constrained such that target and masker words always differed from each other, for any given word position in the target and masker sequence. One talker was used for the target, the other for the masker. Prior to concatenation, each utterance was initially time-scaled to a duration of 300 ms (***Hejna and Musicus, 1991***). In addition, 300 ms silences were included between consecutive words, such that the total duration of each target sequence equaled 15 s.

### Vocoding

Next, the target word sequences were vocoded through an analysis-, followed by a synthesis-filtering stage. For the analysis stage, each word sequence was filtered into 16 adjacent spectral bands, with center frequencies from 300 to 10 kHz. These spectral bands were spaced linearly along the cochlea according to Greenwood’s scale, with a distance of more than one equivalent rectangular cochlear bandwidth between neighboring filters (***Greenwood, 1990***; ***Chen et al., 2011***). Analysis filters had a simulated spectral width of 0.37 mm along the cochlea (***Greenwood, 1990***) or approximately 1/10th octave bandwidth, had a 72 dB/octave frequency roll-off and were implemented via time reversal filtering, resulting in zero-phase distortion. In each narrow speech band, the temporal envelope of that band was then extracted using Hilbert transform. Broadband uniformly distributed white noise carriers were multiplied by these envelopes. For the synthesis stage, these amplitude-modulated noises were then processed by the same filters that were used in the analysis stage. Depending on the experimental condition, a subset of these sixteen bands was then added, generating an intelligible, spectrally sparse, vocoded target sequence.

### Target/Masker Configurations

A target sequence was always presented simultaneously with a masker sequence. Analogous to an established behavioral paradigm for assessing IM, we used two different masker configurations, consisting of different-band-speech or different-band-noise (***Arbogast et al., 2002***). In the SPEECH condition, the masker sequence was designed similarly to the target except that it was constrained such that 1) the target and masker words were never equal at the same time and 2) the masker was constructed by adding the remaining seven spectral bands not used to build the target sequence. In the NOISE condition, the masker sequence consisted of 300-ms long narrowband noise bursts that were centered at the seven spectral bands not used to build the target sequence. All processing steps were identical to the SPEECH condition, expect that, instead of being multiplied with the Hilbert envelopes of the masker words, the noise carriers were multiplied by 300-ms long constant-amplitude envelopes that were ramped on and off with the target words (10 ms cosine squared ramps). Figure 1A shows a representative spectral energy profile for a mixture of target (brown) and SPEECH (black) sequences. Note that the spectrum of a mixture of target and NOISE samples comprised of similar frequency bands would look visually indistinguishable from target in SPEECH and is thus not shown here (c.f. ***Arbogast et al.*** (***2002***)).

In experiment 1, target and either a different-band speech or a different-band-noise masker were presented binaurally (Figure 1B). The target had a left-leading interaural time difference (ITD) of −500 *μs*. The masker sequence had a right-leading 500 *μs* ITD, resulting in two possible target/masker configurations, called SPEECH (different-band-speech with 500 *μs* ITD) *vs* NOISE (different-band-noise with 500 *μs* ITD). The target and masker were each presented at 59 dBA, as calibrated with a 1-kHz tone that was presented at the same root mean square as the target and masker and recorded with KEMAR microphones (Knowles Electronics model KEMAR 45BB). As a result, the broadband Target-to-masker energy ratio (TMR) equaled 0 dB. However, at each of the center frequencies of the nine vocoded spectral bands that made up the target, the TMR equaled 93 dB or more.

In experiment 2, the masker always consisted of a different-band-speech sequence. Target and masker sequences were presented in two possible configurations. The first configuration was identical to the SPEECH condition of experiment 1, with the target presented binaurally with a −500 *μs* ITD and a SPEECH masker at 500 *μs* ITD. In the second “SPEECH-oppo” configuration, a target and different-band-speech masker were presented to opposite ears, with the target presented monaurally to the left, and a different-band-speech masker monaurally to the right ear (Figure 2).

### Behavioral Task

The auditory task consisted of twelve 45-second long blocks. To familiarize the listener with the target voice, at the beginning of each block, we presented a 3-second long cue sentence with the target talker’s voice and instructed the listeners to direct their attention to this talker. The cue sentence was “Bob found five small cards,” and was processed identically to the target speech for that block (same spectral bands, same binaural configuration). Each block then consisted of a 15-second long acoustic mixture of one randomly generated target and one randomly generated masker sequence, followed by a rest period of 30 seconds of silence. Moreover, at the end of each auditory task block, we added a random silent interval (mean: 3.8 s, variance: 0.23 s, uniform distribution). In experiment 1, we randomly interleaved six SPEECH blocks with six NOISE blocks, whereas in experiment 2, we randomly interleaved six SPEECH blocks with six SPEECH-oppo blocks. The spectral bands of the vocoded target and masker were fixed within each block and randomly interleaved across blocks.

Listeners were instructed to press a button each time the target talker to their left side uttered any of the four color keywords, while ignoring all other words from both the target and the masker. A random number (between three and five) of color words in the target voice would appear during each block. No response feedback was provided to the listener.

### Behavioral Detection Threshold

Throughout each block we counted *N_B_*, the number of intervals that the listener pushed the button of the response interface. If a button push occurred within 200 to 600 ms after the onset of a target keyword, the response was scored as a hit. Absence of any button push response in the same time period was scored as a miss. The observed percent correct was calculated by dividing the number of hits by the total number of target keywords during that block.

The baseline guessing rate was estimated via a bootstrapping analysis that calculated the chance percent correct that a simulated listener would have obtained by randomly pushing a button N times throughout that block. Specifically, to estimate the chance percent of keywords guessed correctly via random button push, for each particular listener and block, we randomly shuffled *N_B_* button push intervals across the duration of that particular block’s target sequence and counted the number of keywords guessed correctly, then repeated the process by randomly shuffling again for a total of 100 repetitions. To correct for bias, the observed *vs* chance percent correct scores were then converted to d’-scores, by calculating the difference in z-scores of observed percent correct *vs* chance percent correct (***Klein, 2001***).

### Behavioral Pilot Control

Behavioral pilot testing established the presence of IM in our stimuli, while also verifying that the high- *vs* low-IM conditions tested *via* fNIRS resulted in comparable speech intelligibility. Inside a double-walled sound-attenuating booth (Industrial Acoustic Company), we tested 11 normal-hearing listeners using the same auditory testing equipment and the same speech detection task that we used during the fNIRS recordings, except that listeners had their eyes open during this pilot testing.

In addition, using vocoded stimuli that were recorded by the same talkers as the stimuli used for the speech detection task, we assessed speech identification thresholds by using the coordinate response measure task (***Brungart, 2001b***; ***Kidd Jr et al., 2008***). Briefly, this task presents listeners with the following sentence structure: “Ready [call sign] go to [color] [number] now.” There were eight possible call signs < Arrow, Baron, Charlie, Eagle, Hopper, Laker, Ringo, Tiger>, the same four colors as in the detection task <red, blue, white, green>, and seven numbers (numbers one through eight, except “seven” because, unlike the other numbers, it consists of two syllables). The target sentence was spoken by the same talker for every trial and always had “Baron” as call sign; the masker was either SPEECH or NOISE from a different talker, and using a different call sign than “Baron.” Listeners were instructed to answer the question “Where did Baron go?” by identifying the color in the target sentence. The masker was held fixed at 65 dB SPL, whereas the target level varied randomly from trial to trial from 45 to 85 dB SPL, resulting in five possible TMRs from −20, −10, 0, 10, and 20 dB. The target levels were randomized such that all five TMRs were tested in random order before all of them were repeated in different random order. Listeners competed 20 trials per TMR, both in SPEECH and in NOISE. In addition, to verify that all listeners could understand the vocoded speech in quiet at the softest target level, prior to testing masked thresholds, listeners completed 20 trials in quiet at 45 dB SPL.

In quiet, all listeners scored at or near ceiling in the identification task (Figure 4A), consistent with previous results that nine-band speech stimuli remain highly intelligible despite vocoding (***Shannon et al., 1995***). Speech identification thresholds were much worse in SPEECH than NOISE thresholds (Figure 4B), confirming that the current stimulus processing produces IM (***Arbogast et al., 2002***). Using Bayesian inference, each listener’s SPEECH and NOISE percent correct speech identification curves were fitted with sigmoidally shaped psychometric functions, as a function of TMR (Matlab toolbox: psignifit; (***Wichmann and Hill, 2001***)). Identification thresholds were defined as the TMR at 50% correct of these fitted functions. Paired t-tests comparing speech identification thresholds between SPEECH and NOISE found that performance was significantly worse in SPEECH [paired t-test, t(10) = 25.4, p<0.001]. The effect size, calculated as the Cohen’s d ratio of the difference in SPEECH and NOISE thresholds divided by the pooled standard deviation across listeners, equaled 4.6. Similarly, speech keyword detectability was better in NOISE than SPEECH, by an average 0.4 d’-units [Figure 4C; paired t-test, t(10)=-2.6, p = 0.027]. Cohen’s d equaled 1.0.

**Figure 4.**
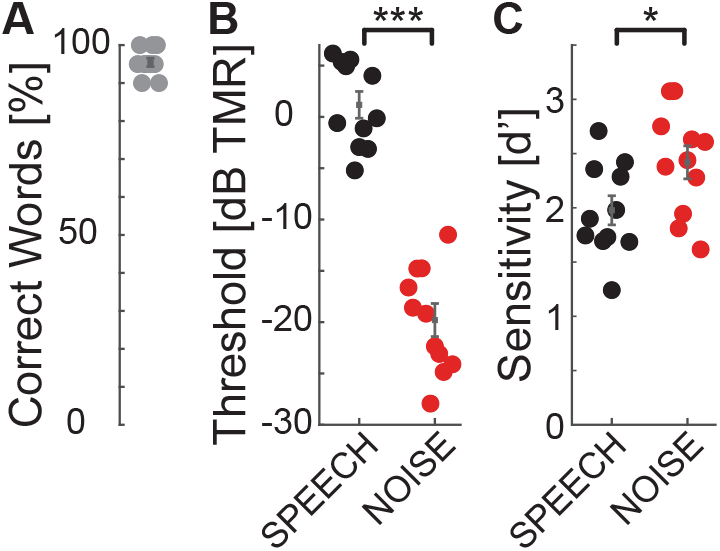
Speech identification and detection performance during pilot testing for SPEECH *vs* NOISE confirm that the SPEECH masker causes IM. The target had a left-leading ITD of −500 *μs*; the masker a right-leading ITD of 500 *μs*. (A) Quiet thresholds. Percent correct keywords identified without masker. (B) Speech identification task. Percent correct keywords identified with SPEECH (black) or NOISE (red) masking. (C) Speech detection task. Sensitivity to keywords with SPEECH (black) or NOISE (red) masking.

We wished to eliminate the possibility of artifacts from eye movements and visual attention in our hemodynamic traces. Moreover, we wished to have comparable task difficulty across the tested conditions with fNIRS. Therefore, we next selected the keyword detection task for neuroimaging, because listeners could perform it with minimal body movement and closed eyes. Moreover, task performance was more comparable across maskers for speech detection *vs* the identification task.

### Neuroimaging Procedure

For both experiments, each listener completed one session of behavioral testing while we simultaneously recorded bilateral hemodynamic traces in the vicinity of STG and cIFS, using fNIRS. Throughout testing listeners held their eyes closed. Traces were acquired in 23-minute sessions, consisting of 11 blocks of a controlled breathing task (9 minutes), followed by a brief break (ca. 2 minutes) and twelve blocks of auditory assessment (12 minutes). The controlled breathing task was identical to our prior methods (see details in ***Zhang et al.*** (***2018***)). Briefly, the task consisted of eleven 45-second-long blocks. In each block, listeners were instructed to breathe in for 5 seconds, breathe out again for 5 seconds. This breathe-in-breathe-out pattern repeated for 6 times (30 seconds in total) before the listeners were instructed to hold breath for 15 seconds. The hemodynamic traces collected during this task establish a baseline dynamic range, from baseline to saturation, over which the optical recordings could vary for each particular listener, recording day and ROI. The auditory assessment was the behavioral detection task described above (see Behavioral Pilot Control).

### Recording Setup for fNIRS

The listener wore insert earphones (Etymotic Research ER-2) and a custom-made fNIRS head-cap and held a wireless response interface in the lap (Microsoft Xbox 360 Wireless Controller; Figure 1A). Acoustic stimuli were generated on a laptop (Lenovo ThinkPad T440P) with Matlab (Release R2016a, The Mathworks, Inc., Natick, MA, USA), D/A converted with a sound card (Emotiva Stealth DC-1; 16 bit resolution, 44.1 kHz sampling frequency) and presented over the insert earphones. This acoustic setup was calibrated with a 2-cc coupler, 1/2” pressure-field microphone and a sound level meter (Bruel&Kjaer 2250-G4). The testing suite had intermittent background sound level with peak levels of 44 dBA (moderately quiet university hallway with noise from staff walking by). Together with the ER-2 insert earphones, which provide approximately 30 dB attenuation, the effective background noise level reaching the listener’s tympanic membrane was 14 dB A, i.e., moderately quiet.

A camera-based 3D-location tracking and pointer tool system (Brainsight 2.0 software and hardware by Rogue Research Inc., Canada) was used to place the optodes above the left and right cIFS and STG, referenced to standardized brain coordinates (Talairach Atlas; ***Lancaster et al.*** (***2000***)). A custom-built head cap, fitted to the listener’s head via adjustable straps, embedded the optodes and held them in place.

Hemodynamic traces were recorded with a 4-source and 16-detector continuous-wave fNIRS system (690 nm and 830 nm optical wavelengths, 50 Hz sampling frequency; CW6, TechEn Inc). The spatial layout of the optical source-detector pairs was custom-designed to cover each of the four ROIs using cross-wise deep quadruple channels with source-detector distances of 3 cm (solid lines in the bottom insert in Figure 1A) and one short separation channel with a source-detector distance of 1.5 cm (dashed lines in bottom insert of Figure 1A). For each of the resulting 16 deep and 4 shallow source-detector pairs, we then used simulated photon paths to estimate a sensitivity map across the surface of brain by mapping the light paths through a standardized head (Figure 1C, AtlasViewer; (***Aasted et al., 2015***)).

### Signal Processing of the fNIRS traces

Raw fNIRS traces were processed to estimate hemodynamic activation strength (Supplement 2 Figure 1A). We first used HOMER2 to process the raw recordings during both the breath holding and auditory tasks, at each of the 16 deep and four shallow source-detector channels (***Huppert et al., 2009***). Specifically, the raw recordings were band-pass filtered between 0.01 and 0.1 Hz, using time-reversal filtering with a fifth order zero-phase Butterworth filter for high pass filtering and time-reversal filtering with a third order zero-phase Butterworth filter for low pass filtering (commands *filtfilt* and *butter* in Matlab 2016). Next, we removed slow temporal drifts in the band-pass filtered traces by de-trending each trace with a 20th-degree polynomial (***Pei et al., 2007***). To suppress artefacts due to sudden head movement, these de-trended traces were then transformed with Daubechies-2 base wavelet functions. Wavelet coefficients outside the one interquartile range were removed, before the remaining coefficients were inversely transformed (***Molavi and Dumont, 2012***). We then applied a modified Beer-Lambert law to these processed traces, resulting in the estimated oxygenated hemoglobin (HbO) and deoxygenated hemoglobin (HbR) concentrations for each channel (***Cope and Delpy, 1988***; ***Kocsis et al., 2006***). To obtain hemoglobin changes relative to the maximum dynamic recording range for each individual listener and recording site, we then applied a normalization step. Specifically, for each listener and each of the 20 source-detector channels, we divided the HbO and HbR concentration from the task conditions by the peak of the HbO concentration change during the controlled breathing task, resulting in normalized HbO and HbR traces for each channel. Finally, we averaged the four deep channels at each ROI, resulting in a total of four task-evoked raw hemoglobin traces per ROI and listener (deep and shallow, HbO and HbR). We previously found that this dynamic range normalization step helps reduce across-listener variability in our listener population with a diverse range of skin pigmentations, hair consistencies and skull thicknesses (***Zhang et al., 2018***).

### Hemodynamic Activation

To estimate auditory-task-evoked neural activity predicted by fixed effects of high- *vs* low-IM, for each of the two experiments, we next fitted a linear mixed effect model (LMEM) to the pre-processed deep HbO and HbR traces (see Supplement 2 for details on the equations). The LMEM model assumes that three main sources of variance shape the HbO and HbR traces: 1) a task-evoked response with IM independence (significant task-evoked activation that does not covary with IM vulnerability), 2) a task-evoked response with IM dependence (significant task-evoked activation that covaries with IM vulnerability), and 3) nuisance signals, deemed to be unlikely of neural origin. In addition, the LMEM includes the following factors that are known to drive neural response changes in STG and cIFS: audibility as modelled through left and right across-frequency average PTAs, and plasticity as modelled through change in output attributed to block number. To allow direct comparison of the masker evoked responses across different ROIs, all *β*_*i*_ were referenced relative to the SPEECH recordings in left cIFS.

To estimate whether a neural response captures behavioral phenotypes for vulnerability to IM, for each listener, masker configuration and ROI, we calculated the predicted total HbO and HbR responses from the LMEM weights, ignoring nuisance signals, PTA and plasticity. Using the peak height of the reconstructed HbO or HbR traces as a measure of that ROI’s neural recruitment for that masker, we then evaluated whether that ROI’s hemodynamic recruitment correlated with the listener’s behavioral d’ sensitivity to IM.

## Acknowledgements

The authors would like to thank Denis Pelli and Barbara Shinn-Cunningham for inspiring discussions about this work and Tara Lynn Alvarez for giving us access to a NIRS machine. We are grateful to the agencies that funded this work. This work was supported in part by NIH R56-DC017481 and NJACTS UL1TR003017 to A Ihlefeld. The NIRS scanner was supported by NSF MRI CBET 1428425 to TL Alvarez.

## Supplement 1

### Differences between behavioral pilot *vs* fNIRS testing

During behavioral pilot testing, a significant but small effect of masker emerged in the speech detection task. However, during fNIRS testing, any differences between SPEECH *vs* NOISE in the same behavioral task were too small to reach statistical significance. Specifically, averaged across listeners, speech detection performance in SPEECH equaled 1.97 (S.E. 0.11) during pilot testing as compared to 1.26 (S.E. 0.22) in experiment 1 and 1.71 (S.E. 0.16) in experiment 2. Across-listener average speech detection performance in NOISE equaled 2.41 (S.E. 0.13) during pilot testing *vs* 1.56 (S.E. 0.21) in experiment 1.

The acoustic delivery of stimuli was identical for fNIRS testing and the behavioral pilot, except that testing happened in different rooms. The fNIRS testing suite had environmental background sound, but it was modest. Indeed, the energy reaching the ears from environmental sound in the fNIRS suite was 50 dB softer than either the masker or target source, presumably only subtly worsening EM or not at all, as compared to the behavioral pilot suite. This hints that the overall reduced performance during fNIRS testing is due to listeners being either more distracted and/or having to put more effort into performing the behavioral task when wearing fNIRS head caps.

## Supplement 2

### LMEM

For each experiment, listener and source-detector pair, full hemodynamic traces were pre-processed (Supplement 2 Figure 1A) before task-evoked responses were estimated *via* LMEM (Supplment 2 Figure 1B).

**Supplement 2 Figure 1.**
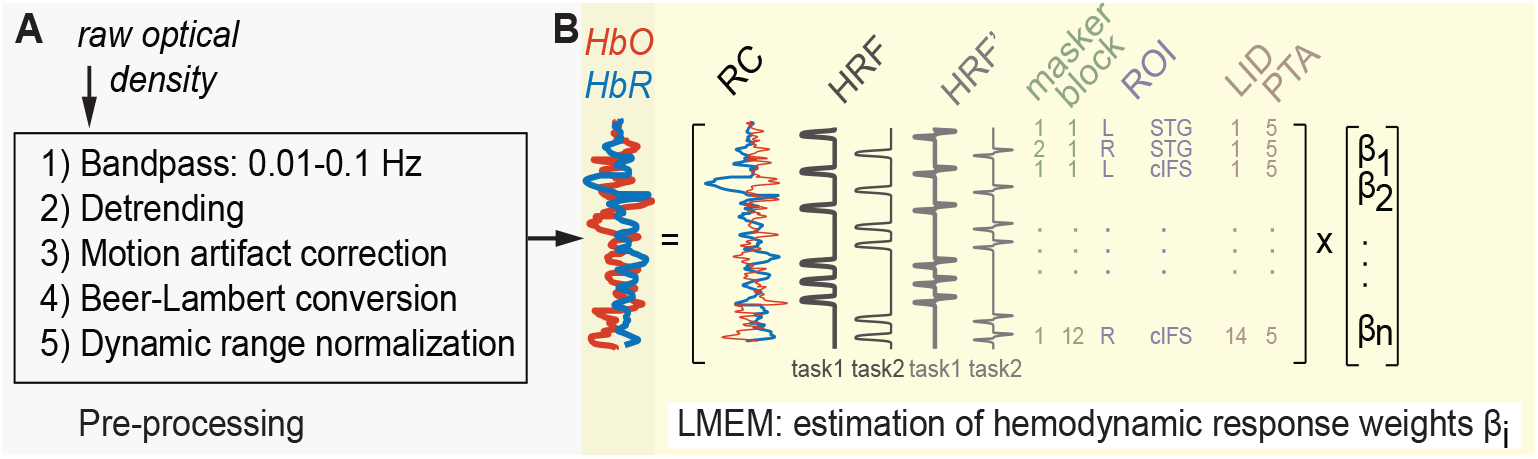
For each experiment, all recorded traces were fit with one LMEM. (A) Signal pre-processing steps. (B) Illustration of default effects in the LMEM.

The hemodynamic response function (HRF, ***Lindquist et al.*** (***2009***) is described by:

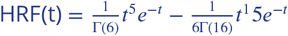

A single LMEM per experiment then fitted these normalized full HbO and HbR traces as follows:

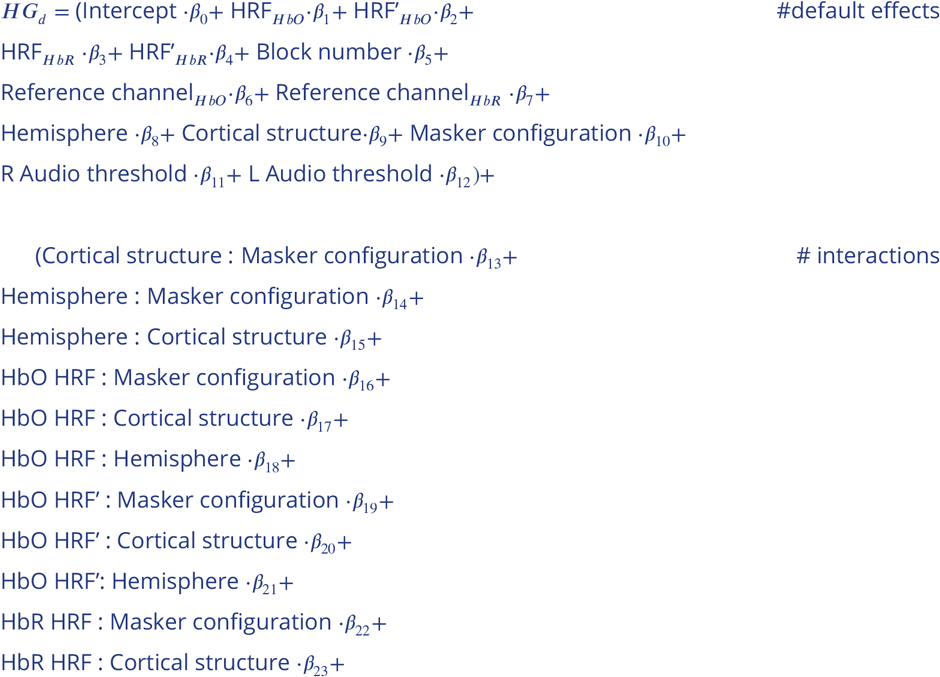

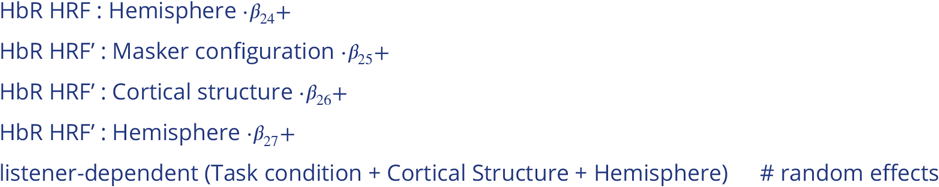

where *HG_d_* is a two-dimensional vector of the normalized full <u>h</u>emo<u>g</u>lobin concentrations, HbO and HbR, recorded from the <u>d</u>eep source-detector channels. The *β_i_* weights associated with each term are a linear measure of how much the term affected the recorded hemoglobin concentration change, relative to the reference condition of SPEECH in left cIFS.

To adjust the onset of the fitted functions to each individual, the LMEM included HRF’, the first derivative of HRF (***Uga et al., 2014***).

Moreover, the LMEM default effects Hemisphere, Cortical structure, and Masker configuration each were two-level categorical variables representing two hemispheres (left *vs* right, *β*_8_), two cortical structures (cIFS *vs* STG, *β*_9_) and two task conditions per experiment (SPEECH *vs* NOISE in experiment 1; SPEECH *vs* SPEECH-OPPO in experiment 2, *β*_10_). Together, these default effects estimated task-evoked responses in the HbO and HbR traces. In addition, the LMEM included factors that are known to drive neural response changes in STG and cIFS: plasticity, modelled through block number (*β*_5_), as well as peripheral hearing, modelled through each individual listener’s across-frequency average left and right PTA (*β*_12_ and *β*_13_). Finally, cardiovascular nuisance signals unlikely to be of neural origin were regressed out via the shallow source-detector Reference channels (RC; *β*_6−7_) in the default model.

As a result, this LMEM implicitly considered that HbR hemodynamic responses are generally much smaller in amplitude and build up more slowly, as compared to HbO (***Watanabe et al., 1996***; ***Sato et al., 2004***). Specifically, HRFs for HbR and HbO were of the same overall canonical functional form. However, to capture potentially different amplitudes and temporal onsets of HbO and HbR, the LMEM fitted HRF and HRF’ amplitudes and their interactions with Masker configuration, Hemisphere and Cortical structure separately for HbO *vs* HbR (*β*_1−4_;(***Niioka et al., 2018***)).

Finally, to regress out idiosynchratic listener-dependent effects on HbO and HbR traces (***Sato et al., 2005***; ***Minati et al., 2011***), the LMEM included random effects for each listener of Masker configuration, Cortical structure and Hemisphere. Note that we initially explored a range of statistical models. We deemed this LMEM model best in terms of explanatory power and parsimony, because it yielded low overall Akaike’s Information Criterion and Bayesian Information Criterion scores (***Anderson and Burnham, 2002***).

## Supplement 3

### Temporal Buildup

Using PET, prior work discovered stronger bilateral STG activation for speech masked by speech relatively to a speech masked by speech baseline (***Scott et al., 2009***), a finding confirmed by the current results via fNIRS. That prior work assessed speech identification while participants listened passively (***Scott et al., 2009***). In contrast, here, hemodynamic responses were recorded while listeners were actively engaged in a speech detection task. Of note, the prior study also showed that the left STG was more strongly recruited than right STG under IM (***Scott et al., 2009***). To examine hemispheric differences, we compared the LMEM predicted hemodynamic response across left and right hemisphere for STG, and, separately for STG. However, for the stimuli tested here during active listening, no robust hemispheric differences in STG activation were revealed.

Furthermore, frontal cortex peak responses during an EM task were found to lag behind STG responses, by approximately 1.5 seconds, when normally-hearing listeners were assessed with fNIRS while listening to vocoded speech in noise (***Wijayasiri et al., 2017***). To analyze the temporal buildup of the task-evoked responses, we subtracted the responses attributed by the LMEM to the STG from those attributed to the cIFS.

**Supplement 3 Figure 1.**
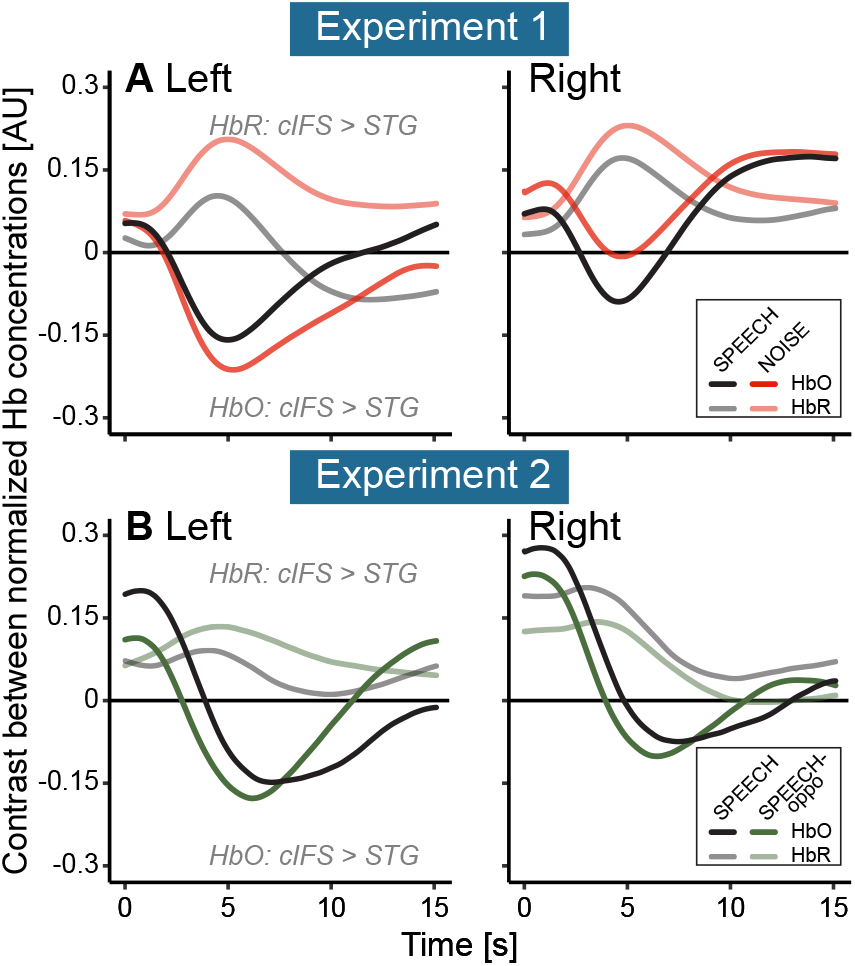
For the first 2-5 seconds of the task, STG was more active than cIFS, whereas cIFS responded more strongly afterwards, with both regions being balanced in their relative activations near the end of the task interval (15 s). This temporal buildup was observed in both hemispheres, for all tested masker configurations, and for both HbO and HbR. Note that HbO (darker lines) and HbR (lighter lines) are anti-correlated, here, as expected. (A) Relative to each ROIs own peak activation levels, STG is slightly more strongly activated than cIFS, for both SPEECH (black) and NOISE in experiment 1(red), in both the Left and the Right hemisphere. (B) The temporal buildup of dominant STG *vs* cIFS activity in experiment 2 are qualitatively comparable to the results in experiment 1 (compare black lines in top *vs* bottom plots. Moreover, the pattern where early STG activity emerges prior to stronger cIFS recruitment also holds for SPEECH-oppo.

In both experiments, masker-evoked differences in overall recruitment of STG *vs* cIFS varied over time. In experiment 1, STG was slightly more strongly recruited during the first 2 seconds of the task interval, before the recruitment between STG and cIFS became more balanced, in both the left and right hemispheres (Supplement 3 Figure 1A). Similarly, in experiment 2, within each hemisphere, STG was relatively more engaged than cIFS during the first 2-5 seconds of the task, followed by stronger HbO and HbR recruitment in the cIFS region (Supplement 3 Figure 1B). Thus, the current temporal buildup results are consistent with prior findings that STG activates before frontal regions, for both EM and IM (***Wijayasiri et al., 2017***).

